# Survey of bacteria associated with western corn rootworm life stages reveals no difference between insects reared in different soils

**DOI:** 10.1101/365296

**Authors:** Dalton C. Ludwick, Aaron C. Ericsson, Lisa N. Meihls, Michelle L.J. Gregory, Deborah L. Finke, Thomas A. Coudron, Bruce E. Hibbard, Kent S. Shelby

**Author notes:** Current Address: USDA-ARS, 2217 Wiltshire Rd., Kearneysville, WV 25430, USA. Current Address: Evogene, Ltd., 1005 N. Warson Rd., Saint Louis, MO 63132, USA. **Author contact information** DCL. **DATA AVAILABILITY STATEMENT** All data are publicly available as Bioproject PRJNA422802, in the NCBI Sequence Read Archive (SRA) database.

## Abstract

Western corn rootworm (*Diabrotica virgifera virgifera* LeConte) is a serious pest of maize (*Zea mays* L.) in North America and parts of Europe. With most of its life cycle spent in the soil feeding on maize root tissues, this insect is likely to encounter and interact with a wide range of soil and rhizosphere microbes. Our knowledge of the role of microbes in pest management and plant health remains incomplete. An important component of an effective pest management strategy is to know which microorganisms are present that could play a role in life history or management. For this study, insects were reared in soils from different locations. Insects were sampled at each life stage to determine the possible core bacteriome. Additionally, soil was sampled at each life stage and resulting bacteria were identified to determine the contribution of soil to the rootworm bacteriome, if any. We analyzed the V4 hypervariable region of bacterial 16S rRNA genes with Illumina MiSeq to survey the different species of bacteria associated with the insects and the soils. The bacterial community associated with insects was significantly different from that in the soil. Some differences appear to exist between insects from non-diapausing and diapausing colonies while no significant differences in community composition existed between the insects reared on different soils. Despite differences in the bacteria present in immature stages and in male and female adults, there is a possible core bacteriome of approximately 16 operational taxonomic units (*i*.*e*., present across all life stages). This research may give insights into how resistance to Bt develops, improved nutrition in artificial rearing systems, and new management strategies.

## Introduction

Several studies have evaluated the microbial communities associated with lepidopteran pests and other insects that attack food crops (1–4). Interestingly, shifts in community composition or absence of bacteria can reduce the effectiveness of widely adopted management tactics such as crop rotation or maize expressing *Bacillus thuringiensis* Berliner (Bt) proteins. However, few studies have been conducted to document microbiomes within beetles attacking crops (5).

The western corn rootworm (*Diabrotica virgifera virgifera* LeConte, WCR) is a chrysomelid beetle whose larvae cause damage to maize root systems. While native to North America, this pest was introduced multiple times to Europe over 20 years ago (6). Most recent estimates indicate this pest causes two billion dollars (USD) in yield loss and control costs worldwide annually (7, 8), and any regions growing maize should monitor for the presence or arrival of this species. Since its discovery as a pest of maize, the primary control tactic has been crop rotation (9). Recently, transgenic maize hybrids expressing insecticidal proteins from Bt have been used to reduce root damage and economic losses. However, both of these control strategies have instances of failure in the United States of America (10–15).

Neonate rootworm larvae (WCR and *D*. *barberi* Smith & Lawrence) burrow through the soil searching for maize root tissues, and then through maize roots while feeding on root tissue. Thus, larvae of these species are exposed to many species of bacteria and fungi in the soil and rhizosphere. The diversity of bacteria encountered is reflected on larval surfaces and digestive tracts. The microbiomes of larvae and later life stages may be assembled from bacterial and fungal species present during larval development in soil.

Insect gut microbiomes are known to influence many aspects of insect growth, nutrition, reproduction, Bt resistance, and pathogen resistance (1, 16–22). Gut microbiota have been shown to affect the response of insects to Bt proteins in Lepidoptera (17, 20–22) and in mosquitoes (23), but this has not been investigated for Coleoptera. In the Old World bollworm (*Helicoverpa armigera* Hübner) the manipulation of the larval gut microbiota with antibiotics results in reduced susceptibility to a commercial formulation of Bt, as well as the purified δ-endotoxins Cry1Ab and Cry1Ac (20). In general, the use of antibiotics to manipulate lepidopteran gut microbiota resulted in reduced mortality due to Bt proteins. Selection experiments with *H*. *armigera* on transgenic plants were also conducted in addition to manipulation of gut microbiota with antibiotics (22). When antibiotics were included, susceptibility to Bt was not altered with increasing generations of selection. However, selection in the absence of antibiotics (gut microbiota unaltered) resulted in a nearly 30% increase in larval survival by the F3 generation (22). Thus, resistance to Bt by *H*. *armigera* developed only when gut microbiota were present. In fact, the reduction in susceptibility to Bt with the addition of antibiotics was greater than the reduction of susceptibility to Bt due to three generations of selection when gut microbiota were present. Gut microbiota were also required for susceptibility of the gypsy moth, *Lymantria dispar* (L.), to Bt proteins (17).

Larval gut tissue of WCR has a diverse microbial community (18, 24). In WCR, a shift in gut microbiota enterotype was associated with increased resistance to soybean defense compounds, which may have contributed to the development of resistance to crop rotation (24). Comparison of gut microbiota between rotation-resistant WCR populations and wild-type WCR populations revealed shifts in the microbial community composition. Manipulation of WCR gut microbiota with antibiotics reduced the resistance to soybean defensive compounds to a level similar to that of wild-type WCR (24).

The contribution of gut microbiota to nutrition, physiology, and Bt resistance in WCR is unknown (18). Feeding of larval WCR on maize root tissue was shown to affect root rhizosphere microbiota composition, indicating a complex, multitrophic interaction (19). Since gut microbiota play a role in Bt susceptibility in lepidopteran pests and a role in crop rotation resistance in WCR, it is reasonable to hypothesize that the microbiota of WCR can affect how larvae respond to Bt toxins expressed in maize. Consequently, a better understanding of which microbes are associated with WCR and how the insects acquire the microbiome is needed. In this study, we focused on the bacteriome. We compared the bacterial composition of WCR grown in two different soils, at each developmental stage, and alongside the soil from which the various life stages were collected and show that WCR larvae can carry particular species across all life stages (*i*.*e.*, a core bacteriome) regardless of the environment.

## Results and Discussion

We conducted the first survey of the bacteriome of WCR and the soil they are found in across all life stages. We investigated the effect of soil origin on the insect bacteriome because WCR occurs across a large region in many different soils throughout the United States of America and Europe. Soil was collected from Higginsville, MO, and the soil bacterial background from which insects emerged was compared to autoclaved soil from Columbia, MO. The results show that earlier life stages reared in soils from different locations contained a significantly different assemblage of bacterial species. However, as the insects matured, those differences declined and all life stages of the insects converged to a similar bacteriome.

Sequencing of WCR and soil samples resulted in a mean (± SEM) of 66,759 (± 3,895) and 72,868 (± 5,308) reads per sample, respectively. To account for the potential influence of differential coverage on downstream analyses, data were randomly subsampled to a uniform depth of 10,000 reads per sample and all subsequent analyses were performed on this rarefied dataset.

Annotated to the taxonomic level of class, the WCR samples were dominated by *Alphaproteobacteria* and *Gammaproteobacteria*, with lower and inconsistent relative abundance of *Actinobacteria*, *Cytophaga*, *Sphingobacteria*, *Betaproteobacteria*, and in the case of surface-sterilized eggs, *Flavobacteriia* and *Deltaproteobacteria* (Fig 1A). Soil samples demonstrated a seemingly more complex composition comprising a greater number of classes and a more even distribution (Fig 1B).

**Fig 1.**
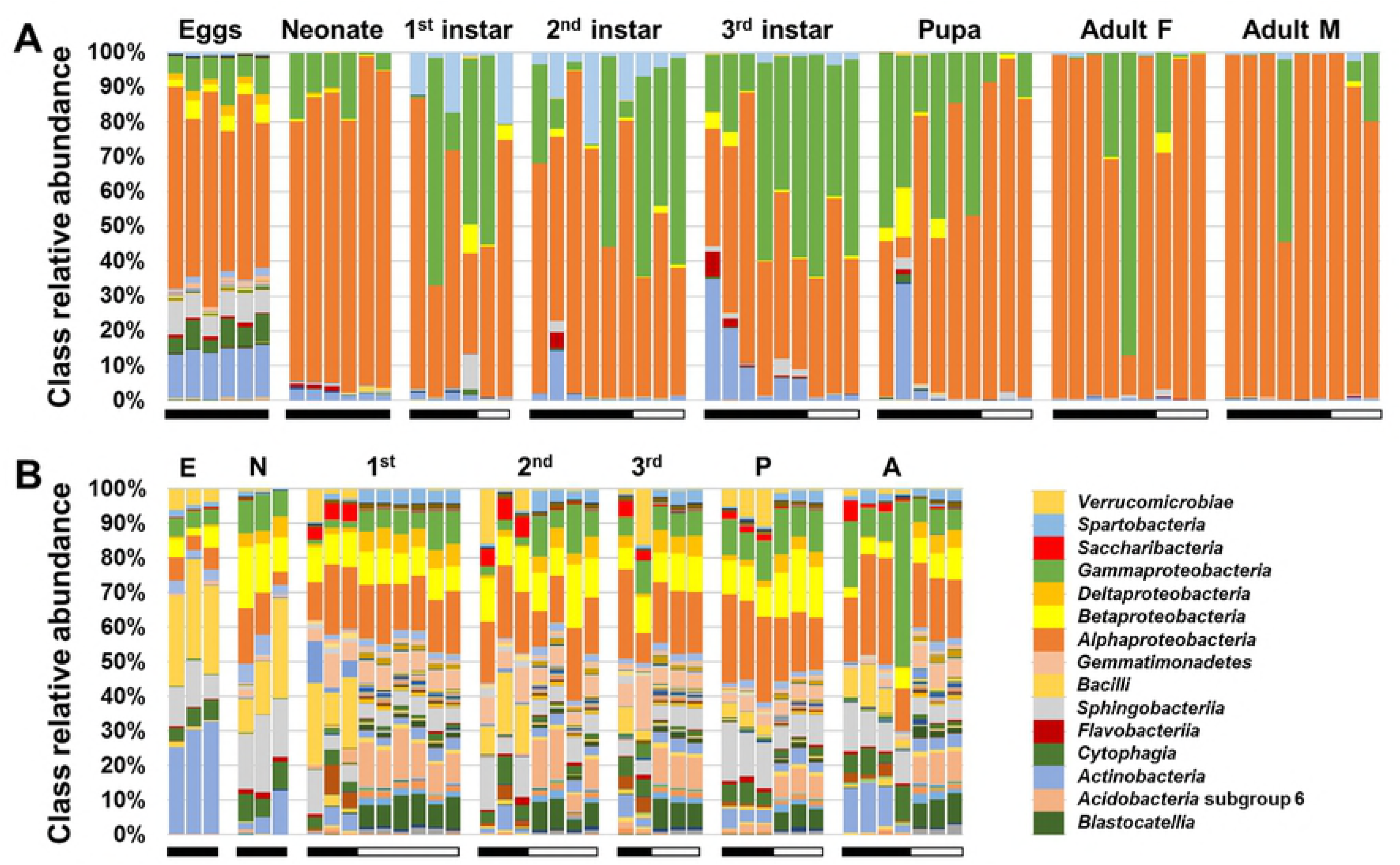
Stacked bar charts showing relative abundances of bacterial classes detected in corn rootworms at different life stages (A) and in soil from which rootworm samples were collected (B). Horizontal bars below the vertical bars indicate original of soil; black bars = Columbia, MO, white bars = Higginsville, MO.

Microbial richness and diversity are often correlated with the health of an ecosystem, be it environmental or host-associated. Richness simply denotes the overall number of detected phylotypes in a sample, whereas Shannon and Simpson diversity indices integrate both the richness and evenness of the distribution of phylotypes in a sample. The underlying assumption is that increased numbers of different taxa and more even distributions of those taxa are representative of ecosystems fostering cross-feeding and syntrophic relationships among microbes. In contrast, low richness or asymmetrical distributions might represent an environment with high selective pressures or the presence of dominant taxa in a competitive environment.

Analyses of richness and diversity of bacterial communities in WCR and in the soil in which they were maintained revealed several interesting trends. To first determine whether the site of soil origin influenced richness, Shannon diversity index, or Simpson diversity index of WCR bacteria, a two-way ANOVA was performed with soil site (*i*.*e*., Columbia or Higginsville) and insect life-stage as fixed variables. Significant main effects of WCR life-stage were detected for richness (*p* < 0.001, F = 8.14), Shannon index (*p* = 0.011, F = 3.48), and Simpson index (*p* < 0.001, F = 5.78). No differences were detected between soil sites for richness, Shannon index, or Simpson index of WCR-associated bacteria (*p* = 0.338, 0.072, and 0.244, respectively). Of note however, similar testing of the soil communities from each site revealed significant site-dependent differences in richness, Shannon index, and Simpson index (*p* < 0.001 for all three metrics, F = 38.52, 197.64, and 25.04, respectively). No life-stage-dependent differences in bacterial richness were detected between the two soil sites, although diversity within soil did significantly vary among life-stages (*p* = 0.030, F = 2.88 and *p* < 0.001, F = 5.53 for Shannon and Simpson indices, respectively).

Collectively, we interpret these data as evidence that the environment has a limited effect on the relative uniformity and richness of the WCR bacteriome. This hypothesis is supported by the nearly log-fold difference in richness between soil and rootworm samples. The fact that no soil-dependent differences were detected in the bacteriome of rootworms themselves, despite the stark differences in the bacterial richness of their respective environments, stands in contrast to the life-stage-dependent differences in richness observed only in the rootworms and not in the soil samples.

Considering WCR samples from the two soils collectively, there was a general trend toward increasing richness in each successive life-stage from egg to pupa followed by a precipitous decline during the pupal molt to adulthood (Fig 2A). Pairwise comparisons of richness between life-stages detected significantly decreased richness of phylotypes in adult WCR relative to several earlier life-stages. Interestingly, an inverse trend was observed in the richness of bacteria in soil samples across life-stages (Fig 2B). In contrast, diversity as assessed via the Simpson index, was higher in sterilized eggs relative to other life-stages while diversity in adult rootworms was much lower (S1A Fig), likely reflecting the increasingly skewed bacterial community structure as the rootworms mature. No life-stage-dependent differences were detected in the diversity of the soil bacterial community (S1B Fig).

**Fig 2.**
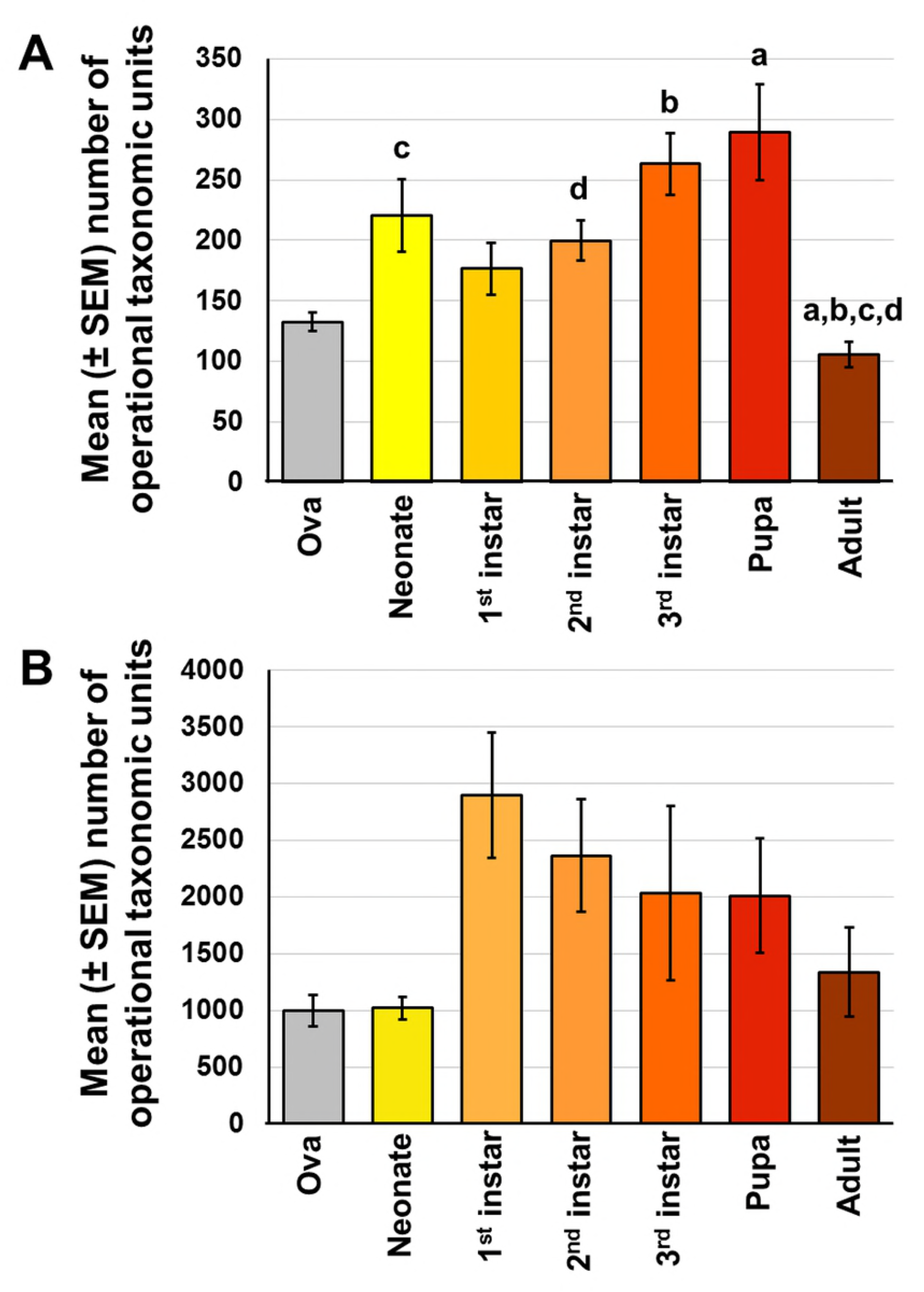
Main effect of life stage on bacterial richness in western corn rootworm (**A**, *p*<0.001), or the soil from which WCR samples were collected (**B**, *p* = 0.040). Significant pairwise differences are indicated by like letters (Kruskal-Wallis one-way ANOVA on ranks with Dunn’s *post hoc*).

In order to provide a more comprehensive comparison of the bacterial communities present in each sample, incorporating not just the number but also the identities of shared and unique taxa, principal coordinate analysis (PCoA) and permutational multivariate analyses of variance (PERMANOVA) were performed to visualize and statistically test for differences in community structure, respectively. With both methods, the similarity of any given pair of samples can be determined several different ways. To ensure that any differences detected were robust and to determine the nature of detected differences, we compared samples using both the Bray-Curtis and Jaccard similarity indices. While the Jaccard index is relatively unweighted and determines sample similarity based on the shared presence or absence of taxa, the Bray-Curtis index is weighted to also incorporate the relative abundance of any shared taxa.

Regardless of the index used, robust compositional differences were detected among all groups with the exception of the WCR samples reared in soil from different sites, again suggesting selection for a specific bacterial community within the rootworms. Specifically, testing for differences using the Bray-Curtis distances detected significant compositional differences between all pairwise comparisons except between WCR samples reared in different soil (Table 1). Accordingly, PCoA demonstrated a clear separation of soil and WCR samples along PC1 (38.1% of the total variation in the dataset), complete separation of soil communities from the two soil sites along PC2, and partial overlap between WCR communities (Fig 3). Testing based on the Jaccard index found significant differences between all pairwise comparisons. Ordination resulted in a similar pattern and the F value generated from the comparison of WCR reared on soil from the two sites was extremely low relative to the other comparisons, despite having the highest total number of samples included in the comparison (Table 2). Collectively, these data complement the analyses of richness and diversity in supporting the hypothesis that WCR select for a limited subset of host-associated bacteria, largely irrespective of their environment.

**Table 1.** Results of PERMANOVA testing for differences in β-diversity between western corn rootworm (WCR) and soil samples collected from two different sites, based on the Bray-Curtis distance. *p* values and F values are shown in the upper right and lower left portions of the table, respectively.

| F values |  | *p* values |  | Soil origin |  | WCR from “X” soil |  |
| --- | --- | --- | --- | --- | --- | --- | --- |
|  |  |  |  | Columbia | Higginsville | Columbia | Higginsville |
| Soil origin | Columbia |  |  |  | 0.0001 | 0.0001 | 0.0001 |
|  | Higginsville |  |  | 27.62 |  | 0.0001 | 0.0001 |
| WCR from “X” soil | Columbia |  |  | 57.08 | 104.5 |  | 0.1498 |
|  | Higginsville |  |  | 38.43 | 119.7 | 1.657 |  |

**Table 2.**
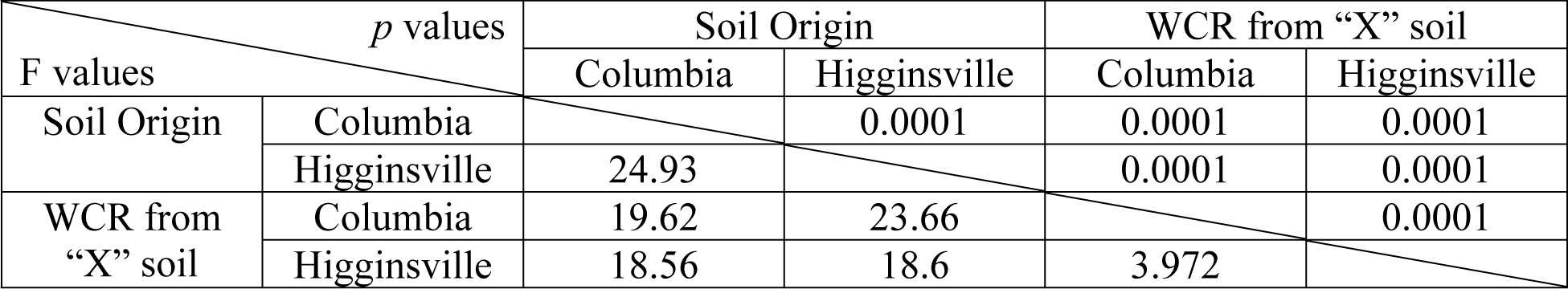
Results of PERMANOVA testing for differences in β-diversity between western corn rootworm (WCR) and soil samples collected from two different sites, based on the Jaccard distance. *p* values and F values are shown in the upper right and lower left portions of the table, respectively.

**Fig 3.**
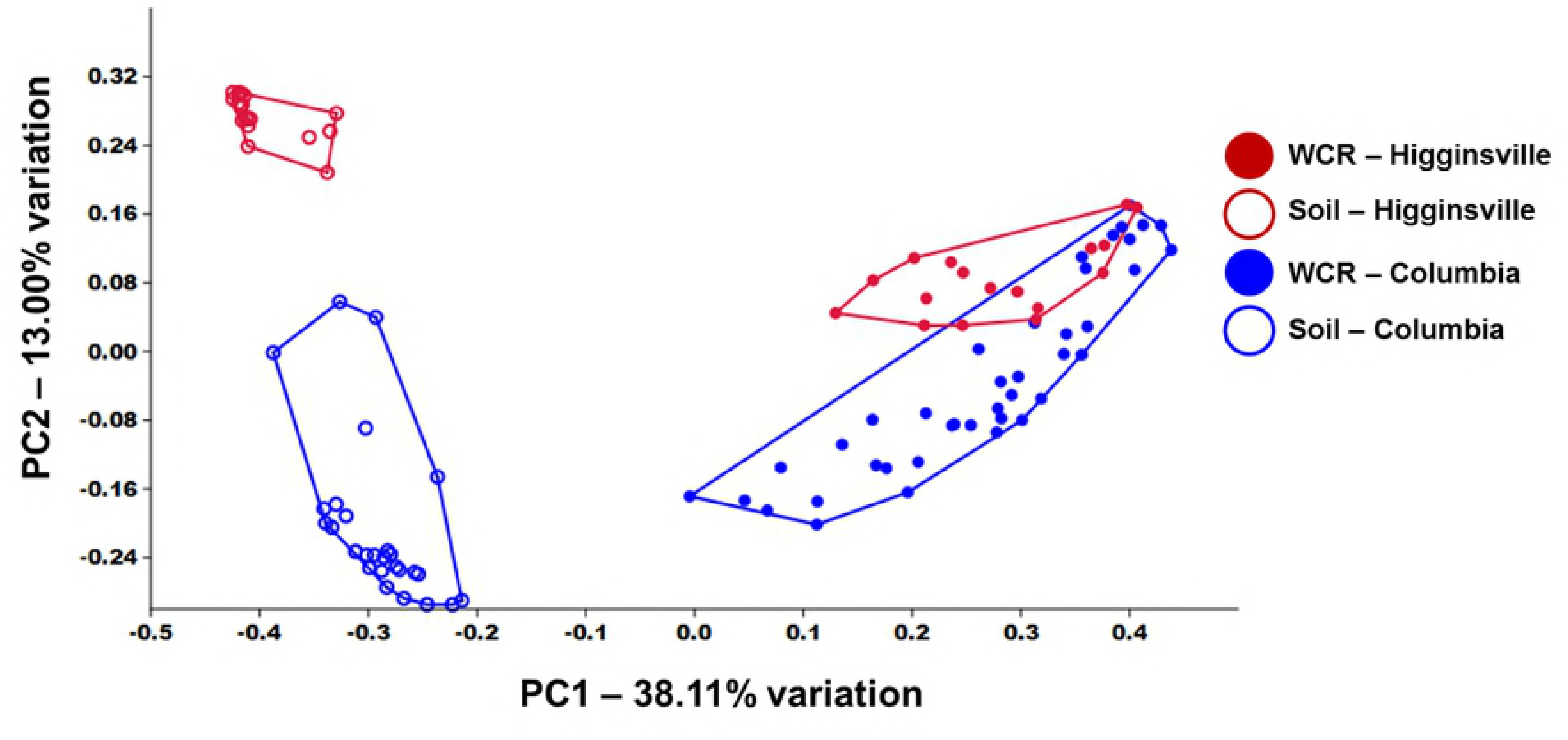
Principal coordinate analysis based on Bray-Curtis similarity between bacterial communities detected in western corn rootworm (WCR) at various life stages and soil samples collected from two different sites.

Annotated to the level of operational taxonomic unit (*i*.*e*., the best taxonomic resolution afforded by the 16S rRNA amplicons), the bacterial composition of the adult WCR was incredibly sparse. Of the 474 operational taxonomic units (OTUs) detected in anywhere from one of 18 (5.6%) to 15 of 18 (83.3%) of the adult rootworms, the mean relative abundance was uniformly below 0.3% (S2 Fig). Conversely, the 13 OTUs detected in 16 or greater of the 18 adult rootworms were present at a mean relative abundance of greater than 1.5%. Notably, 95.4% of the bacterial DNA recovered from adult rootworms was annotated to three OTUs: *Wolbachia* sp. (85.5 ± 24.0% in 18 of 18 adults), unclassified family *Enterobacteriaceae* (6.2 ± 13.0% in 16 of 18 adults), and *Acinetobacter* sp. (4.7 ± 11.6% in 17 of 18 adults).

To determine whether inherent differences exist in the bacteriome of WCR based on genetic background, insects from a colony of wild-type WCR that undergo diapause and an experimental non-diapausing WCR laboratory colony were reared to each life stage in autoclaved soil from Columbia, MO, as previously mentioned. All life stages and corresponding soil samples were collected and processed to extract and purify DNA. The V4 region of the 16S ribosomal gene was amplified and sequenced to putatively identify bacteria.

Once the identities of the bacteria were determined, we compared the bacteriomes between the two colonies using PERMANOVA with Bray-Curtis and Jaccard indices. The two indices revealed different patterns. No significant differences were detected between these colonies with the Bray-Curtis index (*p*=0.10; F=1.90), indicating that insects that do not undergo diapause retain a similar bacteriome as insects that do undergo diapause despite hundreds of generations of laboratory selection. However, PERMANOVA with a Jaccard index revealed significant differences in bacterial communities between insects from a diapausing colony and those from a non-diapausing colony (*p*=0.0001; F=2.90; Fig 4). Insects from both colonies appear to share many dominant taxa while rarer species appear to be isolated to individual colonies.

**Fig 4.**
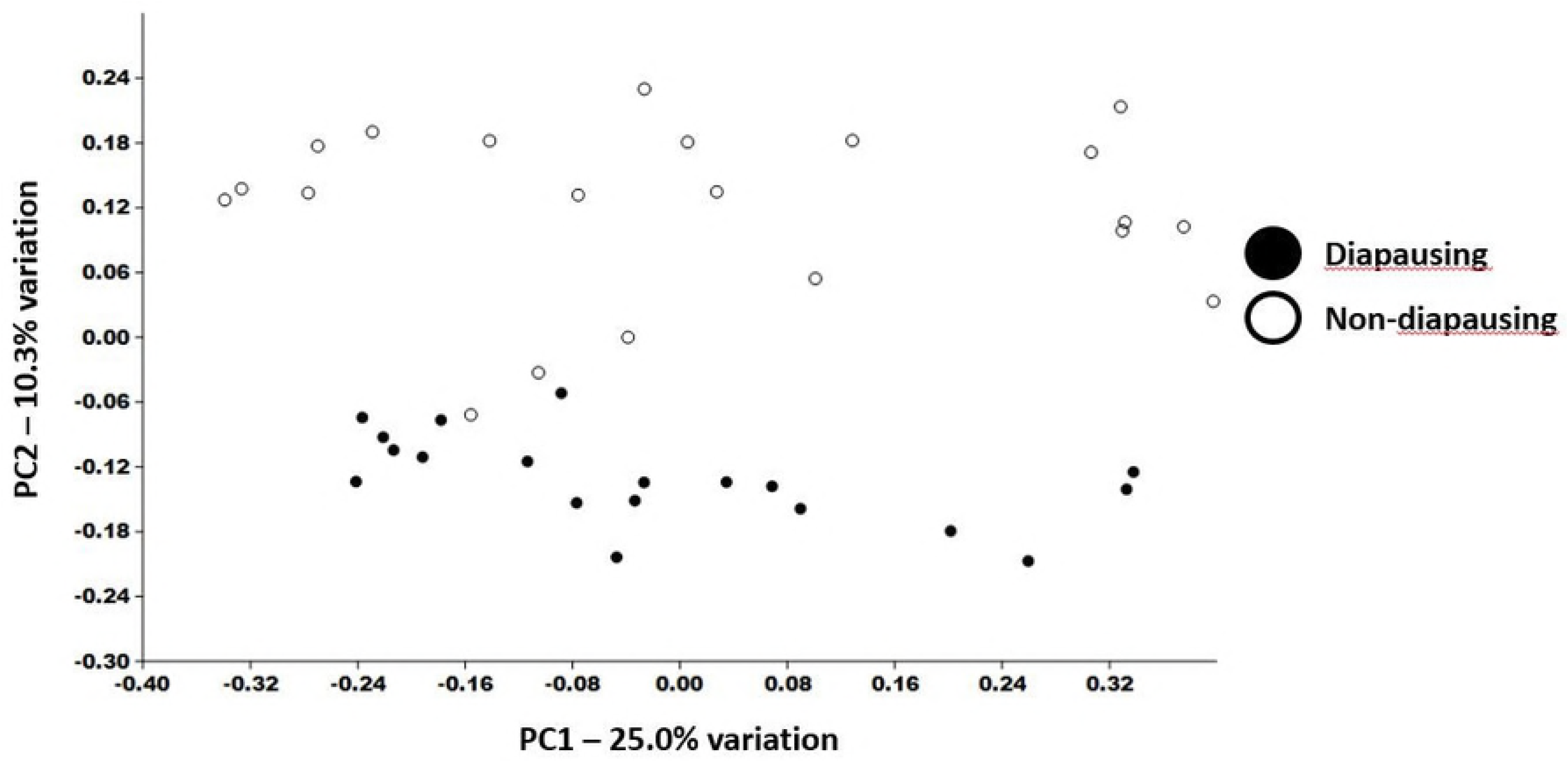
Principal coordinate analysis based on Bray-Curtis similarity between bacterial communities detected in western corn rootworm (WCR) from diapausing and non-diapausing colonies including all life stages, except sterilized ova.

Exploratory studies documenting the bacterial communities in different organisms may lead to new insights as to the role(s) they may fill or even new management tactics. Over 2,200 unique operational taxonomic units (OTUs) were putatively identified in soil and insect samples from both colonies and soils. Our study documented more than 1,100 OTUs present throughout the WCR life cycle. Of these OTUs, 16 were found in every life stage of insects regardless of the colony or rearing soil. We speculate that these 16 OTUs comprise the core bacteriome for WCR. Furthermore, some of these bacteria were never found in the soil suggesting vertical transmission (*i*.*e*., parent to progeny) of bacteria is the most likely mechanism for at least some of the WCR bacteriome (Table 3).

**Table 3.**
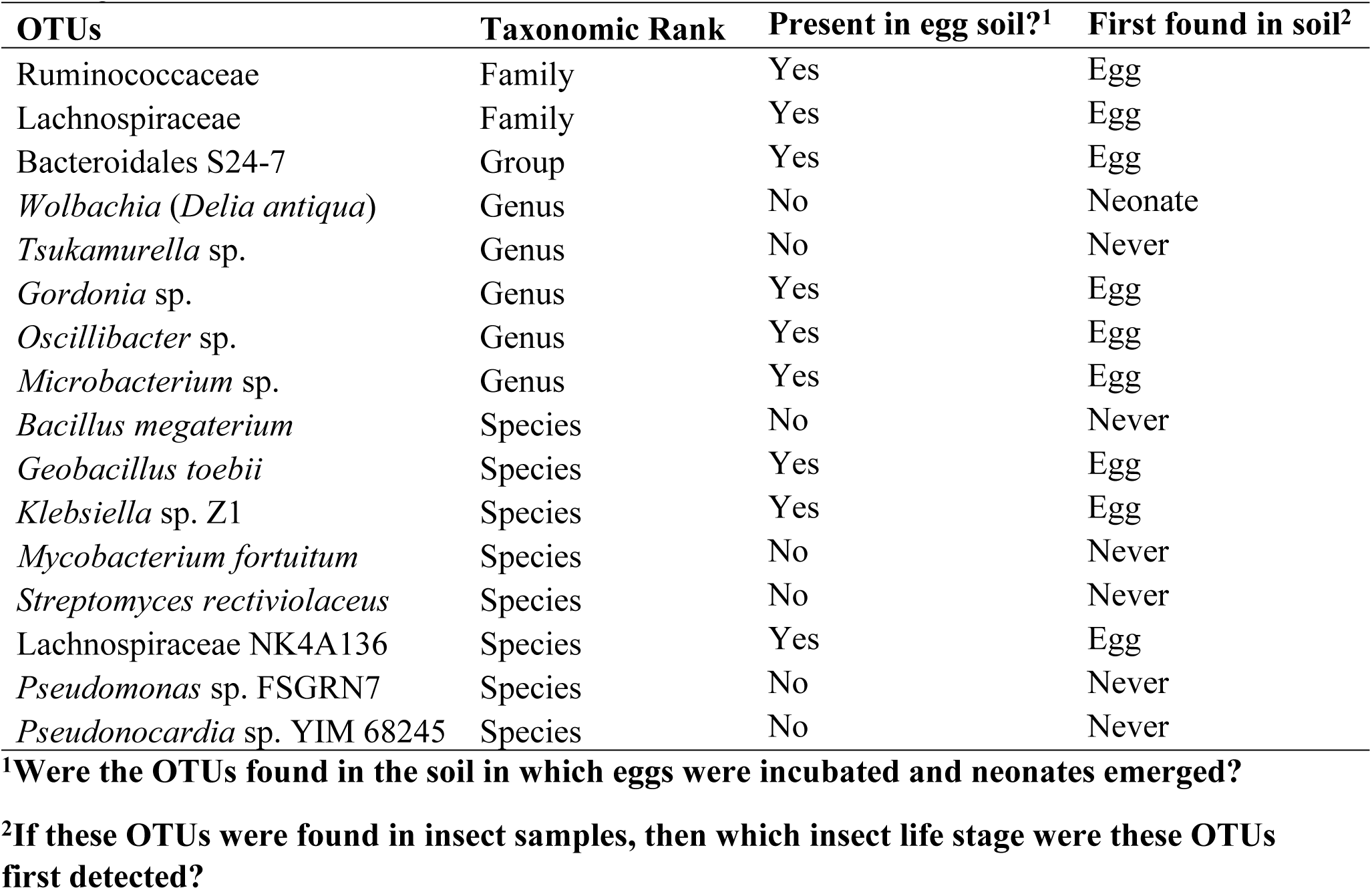
Unique operational taxonomic units (OTUs) found in all insect samples regardless of soil origin.

| OTUs | Taxonomic Rank | Present in egg soil?<sup>1</sup> | First found in soil<sup>2</sup> |
| --- | --- | --- | --- |
| Ruminococcaceae | Family | Yes | Egg |
| Lachnospiraceae | Family | Yes | Egg |
| Bacteroidales S24-7 | Group | Yes | Egg |
| *Wolbachia (Delia antiqua)* | Genus | No | Neonate |
| *Tsukamurella* sp. | Genus | No | Never |
| *Gordonia* sp. | Genus | Yes | Egg |
| *Oscillibacter* sp. | Genus | Yes | Egg |
| *Microbacterium* sp. | Genus | Yes | Egg |
| *Bacillus megaterium* | Species | No | Never |
| *Geobacillus toebii* | Species | Yes | Egg |
| *Klebsiella* sp. Z1 | Species | Yes | Egg |
| *Mycobacterium fortuitum* | Species | No | Never |
| *Streptomyces rectiviolaceus* | Species | No | Never |
| Lachnospiraceae NK4A136 | Species | Yes | Egg |
| *Pseudomonas* sp. FSGRN7 | Species | No | Never |
| *Pseudonocardia* sp. YIM 68245 | Species | No | Never |

<sup>1</sup>Were the OTUs found in the soil in which eggs were incubated and neonates emerged?
<sup>2</sup>If these OTUs were found in insect samples, then which insect life stage were these OTUs first detected?

Many OTUs were discovered in the sterilized eggs of insects from the diapausing colony. However, we cannot be certain whether these bacteria were alive inside the egg or dead on the surface of the egg shell. Given the sculpturing of the chorion, it is possible dead bacteria remained on the surface served as a source of non-viable DNA (19, 25). The protocol we used does not discern between live and dead bacteria. If the bacteria were alive, then it is possible the eggs serve as a source of bacteria that colonize the neonatal gut. There is evidence that some of the bacteria are passed from parents to offspring (Table 3), but we cannot be certain without additional studies. Future experiments should extract rRNA and generate cDNA before sequencing the resulting strands. This method would reduce the likelihood of dead bacterial sequences entering the analysis as RNA degrades rapidly while DNA can persist for many years.

We infer that some of these bacteria may be endosymbionts of WCR as particular OTUs never appeared outside of insect samples (Table 3). However, we used laboratory colonies to make inferences about wild-type populations. In theory, the differences between wild-type populations and laboratory colonies should be minimal. In reality, we simply do not know. The geographic distribution of this insect encompasses most of the United States of America and parts of Europe. The soils across these regions are also diverse as are the management tactics employed by farmers. It stands to reason that the bacterial communities are different within and between fields. Future studies will need to include more samples, samples from different locations across the Corn Belt and other regions, and wild-type specimens to validate or invalidate the findings of this research.

WCR continues to evolve and adapt to the different management tactics that maize growers are implementing now. Future technologies for pest control, including RNA interference, are still years away from field implementation. New tools and knowledge are needed to combat this pest. This study documents the plethora of bacteria encountered by WCR in different soils and identifies a small core bacteriome retained by WCR. Clearly, there is much to learn about the functions of these different bacteria with regards to WCR.

## Materials and Methods

### Insect rearing

Eggs from non-diapausing and diapausing colonies of WCR were obtained from the Agricultural Research Service of the United States Department of Agriculture (USDA-ARS). The non-diapausing colony was derived from the primary non-diapausing colony held at Brookings, SD (26). The diapausing colony eggs were from the primary diapausing colony (27) also held at Brookings, SD, and remained in cold storage until needed.

For the non-diapausing colony maintained in Columbia, MO, adults of both sexes were placed in cages (30×30×30 cm, Megaview Science Co., Ltd., Taichung, Taiwan) with a photoperiod of 14:10 (L:D) h at 25 °C. Adults were supplied with corn leaf tissue, slices of zucchini, an agar gel to serve as a water source, and an artificial diet for adult rearing (Frontier Agricultural Sciences, Newark, DE). Petri dishes with 70 mesh sieved field soil from Columbia, MO, served as an oviposition site for females. The oviposition site was moistened throughout the week and replaced weekly. The eggs in the Petri dish were separated from the soil by washing through a 60 mesh sieve. The eggs were then divided and placed in two plastic containers (15 × 10 cm, GladWare^®^, The Glad Products Company, Oakland, CA) with 70 mesh sieved Columbia, MO, field soil. The plastic containers were covered with lids and placed on the bottom racks of a Percival incubator set to run at 25 °C.

### Seedling Mats

#### Insects from non-diapausing colony

Fifteen seedling mats were planted in March 2016. Each seedling mat contained approximately 15 g of maize seed (Monsanto Company, variety DKC 61-79), 6 cm of autoclaved growth medium, and 80 ml of tap water in a 15 × 10 cm plastic container. The growth medium consisted of a mixture of field soil:Pro-Mix BX potting medium (Premier Horticulture Inc., Quakertown, PA) at a 2:1 ratio (v/v) prior to being autoclaved. Seedling mats were allowed to germinate, and coleoptiles emerged through the soil surface prior to infestation.

Seedling mat containers were placed on the top rack of the same Percival incubator in which eggs were incubated. Data were collected at six time points: 0 d (neonate larvae), 5 d, 10 d, 15 d, 22 d, and adult emergence. Three replicates of each time point were used for this survey. Seedling mats in each replicate were randomly assigned a time point with each seedling mat receiving 25 neonate larvae. The 0 d time point did not require insect feeding and so for this treatment, rather than using seedling mats, 10 neonate larvae were collected directly into 1.5ml microcentrifuge tubes (USA Scientific) and then stored at −80 °C (So-Low, Environmental Equipment, Cincinnati, OH).

For the adult emergence time point in this survey, we planted new maize seeds into a larger container (33 × 19 cm, Sterilite Corporation, Birmingham, AL) and allowed the maize to grow for one week prior to infestation. The first and smaller seedling mat had plant tissue removed before being inverted onto the second and larger seedling mat containing soil from the same site. After one week, the larger seedling mat was covered with a mesh screen to prevent escape of emerging adults.

#### Insects from a diapausing colony

A total of five replications were conducted for this survey. During this survey, two different growth media were used. The first growth medium remained the same as the previous insect survey, while the second growth media was soil collected from a continuous corn field in Higginsville, MO, in July 2016. This soil was not autoclaved and remained enclosed in a metal container until use in October 2016. In addition to the time points listed previously, two types of eggs were sampled: eggs washed from sieved soil, and eggs washed from sieved soil that were then surface sterilized (28).

Once the desired time point was reached, the seedling mats were processed in the same manner as (29). For the 5, 10, 15, and 22 d time points, all aboveground plant material was removed from the container. Next, the soil and root tissue were placed into a Berlese funnel with an attached jar. The jar with a moist filter paper at the bottom was used to collect the larvae. Specimens of each age were transferred from the jar to a microcentrifuge tube at least once every three hours throughout a typical work day. This tube was then immediately placed into the −80 °C freezer for storage until DNA extraction occurred. A new tube was used for each collection time and sample to prevent additional freezing and thawing. During time points when larvae were sampled, soil was also collected from the bottom of the seedling mat prior to drying.

No secondary container was used for the diapausing insect survey, but mesh screens were used to keep the adults from escaping the container. Adult emergence containers were checked daily, and adults from each container on a given day were placed into microcentrifuge tubes. Soil was collected from the soil surface where adults must pass to emerge through the soil.

#### DNA Extraction and Quantification

Whole insects (1-8 larvae/treatment; 1-2 pupae/treatment; a single adult/treatment) were pooled, and DNA extracted using accepted methods (30). The samples were extracted using PowerFecal^®^ DNA Isolation Kit (MO BIO Laboratories, Inc. Catalog No. 12830-50) following the manufacturer’s protocol (https://mobio.com/media/wysiwyg/pdfs/protocols/12830.pdf) with the following modifications: one sterile 0.5 cm diameter stainless steel ball bearing was added to the Dry Bead Tube for each adult and soil sample prior to shaking; shaking time was reduced to 5 minutes for adults and 3 minutes for all other samples. DNA quality and concentration was determined for each sample by Nanodrop 2000 Spectrophotometer (Thermo Scientific, Wilmington, DE) and stored at –80°C.

#### Library construction and sequencing

All PCR and sequencing was performed at the University of Missouri DNA Core. DNA concentration was determined fluorometrically (Qubit 2.0, Life Technologies) prior to analysis. Based on results of fluorometry, all samples were normalized to a standard concentration for PCR amplification. Bacterial 16S rRNA amplicons were generated via amplification of the V4 hypervariable region of the 16S rRNA gene using single-indexed universal primers (U515F/806R) flanked by Illumina standard adapter sequences and the following parameters: 98 °C^(3:00)^+[98°C^(0:15)^+50°C^(0:30)^+72°C^(0:30)^] × 25 cycles +72°C^(7:00)^. Amplicons were then pooled for sequencing using the Illumina MiSeq platform and V2 chemistry with 2×250 bp paired-end reads, as previously described (31).

#### Informatics analysis

All informatics analyses were performed as previously described (32), at the University of Missouri Informatics Research Core Facility. Input is typically for 2×350 bp reads from one of the two MiSeq machines in the DNA Core. The read pairs are joined into contigs by the program FLASH (http://bioinformatics.oxfordiournals.org/content/27/21/2957.long) (33), and culled if found to be short after trimming for a base quality less than 31, and those that are not joined, or are too long or short after contig formation, leaving those that are 275 to 300 nts. Cutadapt (http://iournal.embnet.org/index.php/embnetiournal/article/view/200/479) was used to find and trim the primers from the 5’ and the 3’ ends, culling those contigs lacking both primers. Contigs with the expected number of errors greater than 0.5 were removed by Usearch (http://drive5.com/index.htm), and the remainder were trimmed to length 248. The contig read ids were modified so that samples could be followed throughout by using the Qiime script split_libraries_fastq.py. All samples were then pooled into one FASTA file and metrics for all samples collated into one table. Contigs were clustered *de novo* into an OTU table using the uparse (http://drive5.com/uparse/) algorithm. *De novo* and reference-based chimera detection and removal was performed using Qiime v1.8 (34) software, and remaining contiguous sequences were assigned to operational taxonomic units (OTUs) via *de novo* OTU clustering and a criterion of 97% nucleotide identity. Annotation of selected OTUs was performed using BLAST (35) against the Silva database (https://www.arb-silva.de/) (36) of 16S rRNA sequences and taxonomy. Principal coordinate analysis and PERMANOVA testing were performed using ¼ root-transformed and non-transformed OTU relative abundance data, respectively, using Past 3.16 (https://folk.uio.no/ohammer/past/) (37). Richness, Shannon diversity index, and Simpson diversity metrics were determined in Past 3.16 using Qiime-generated otu_biom.table files.

#### Statistical analysis

Differences in raw and binned OTU richness were tested via ANOVA using SigmaPlot 12.3 (Systat Software Inc., San Jose, CA); *p* values less than 0.05 were considered significant. Differences in the overall composition of the different regions were tested via two- and one-way PERMANOVA of ranked Bray-Curtis or Jaccard distances using the open access Past 3.16 software package (38), downloaded on April 2, 2016.

## Acknowledgements

Sequencing services were performed at the University of Missouri DNA Core Facility by Nathan Bivens and Karen Bromert. We thank Bill Spollen and Christopher Bottoms (University of Missouri Informatics Core), Rebecca Dorfmeyer and Giedre Turner (University of Missouri Metagenomics Center), and Karen Clifford (University of Missouri College of Veterinary Medicine), Julie Barry (USDA-ARS) for insect rearing, Adriano Pereira for assistance with collecting insects, and Chad Nielson and Wade French (USDA-ARS) for maintaining colonies used in this study. Mention of trade names or commercial products in this publication is solely for the purpose of providing specific information and does not imply recommendation or endorsement by the U.S. Department of Agriculture (USDA). USDA is an equal opportunity provider and employer.

## Author Contributions

B.E.H., L.N.M., T.A.C., and K.S.S. secured funding for the project; D.C.L., L.N.M., T.A.C., B.E.H. and K.S.S. conceived and designed the experiments; D.C.L., L.N.M., M.L.G., and A.C.E. performed the experiments; A.C.E., D.C.L. and K.S.S. analyzed the data; D.C.L., L.N.M., A.C. E. and K.S.S. contributed reagents/materials/analysis tools; and D.C.L., L.N.M., M.L.G, A.C.E., D.L.F., T.A.C., B.E.H., and K.S.S. wrote the paper. All authors read and approved the final version.

## Figure Legends

**S1 Fig**. Main effect of life stage on mean Shannon and Simpson diversity indices in western corn rootworms (**A**,*p* < 0.001), or the soil from which the WCR samples were collected (**B**,*p* = 0.040). Significant pairwise differences indicated like letters (Kruskal-Wallis one-way ANOVA on ranks with Dunn’s *post hoc*).

**S2 Fig**. Principal coordinate analysis based on Jaccard similarity between bacterial communities detected in western corn rootworms (WCR) at various life stages and soil samples collected from two different sites.

**S3 Fig**. Number and mean relative abundance (above bars) of operational taxonomic units (OTUs) detected at increasing prevalence in adult western corn rootworm samples.

**SUPPORTING INFORMATION**

